# Localization of Human UBE3A Isoform 3 is Highly Sensitive to Amino Acid Substitutions at p.Met21 Position

**DOI:** 10.1101/2024.02.05.578859

**Authors:** Annelot C. M. van Esbroeck, Rob F. M. Verhagen, Martina Biagioni, Matteo Fossati, Ben Distel, Ype Elgersma

## Abstract

UBE3A encodes three isoforms of Ubiquitin E3 ligase A, which differ in their N-terminal sequence, abundance, and localization. Recently, three individuals diagnosed with Angelman Syndrome have been described who carry a variant that abrogates the start codon of the predominant nuclear isoform 1 (hUBE3A-Iso1^p.Met1Thr^) and concomitantly results in a missense variant in isoform 3 (hUBE3A-Iso1^p.Met21Thr^), which we previously reported to be nuclear enriched as well.

Here, we studied the effect of the p.Met21Thr variant on hUBE3A-Iso3 localization. Recombinant expression of hUBE3A-Iso3^p.Met21Thr^ in U2-OS and mouse neurons revealed similar UBE3A labelling in the nucleus and cytosol, indicating hUBE3A-Iso3 localization is sensitive to amino acid changes at this position. This finding prompted us to revisit hUBE3A-Iso3 localization, since we previously introduced a p.Met21Ala/p.Met22Ala amino acid substitution in hUBE3A-Iso3 and its mouse orthologue mUBE3A-Iso2 to prevent translation of the shorter hUBE3A-Iso1 and mUBE3A-Iso3 nuclear isoforms. Introduction of silent mutations to disfavour translation of the short UBE3A isoforms enabled us to determine the localization hUBE3A-Iso3 and mUBE3A-Iso2 in the absence of amino acid changes at the p.Met21/p.Met22 position, respectively. Surprisingly, hUBE3A-Iso3 localization shifted from predominant nuclear localization for hUBE3A-Iso3^p.Met21Ala^ to a predominant cytosolic localization of the Kozak optimized hUBE3A-Iso3^KOZAK^, while their mouse orthologues mUBE3A-Iso2^p.Met22Ala^ and mUBE3A-Iso2^KOZAK^ both localized predominantly to the cytosol.

Taken together, these experiments indicate that the localization of human UBE3A-Iso3 is highly sensitive to amino acid substitutions at the p.Met21 position and that variants at this position not only abrogate the translation of hUBE3A-Iso1, but can also change the localization of hUBE3A-Iso3.

## Introduction / Results / Discussion

Genetic variants affecting the maternally inherited *UBE3A* gene (NG_009268.1 [MIM: 601623]) are responsible for the severe neurodevelopmental disorder Angelman Syndrome (AS) [MIM05830]. *UBE3A* encodes three isoforms of Ubiquitin E3 ligase A (UBE3A), which differ in abundance as well as localization, driven by their distinct N-terminal sequences resulting from alternative translation start sites. The two major human UBE3A isoforms (hUBE3A-Iso1 (NP_570853.1), ∼80% of total UBE3A and hUBE3a-Iso3 (NP_570854.1) ∼20% of total UBE3A), are highly conserved throughout evolution and have a 96% sequence identity to the corresponding mouse UBE3A orthologues (mUBE3A-Iso3 (NP_001029134.1) and mUBE3A-Iso2 (NP_035798.2) respectively) (1). We previously reported that the short UBE3A isoforms (hUBE3a-Iso1, mUBE3a-Iso3) showed strong nuclear enrichment (1,2). In contrast, the long mouse isoform mUBE3A-Iso2 localized predominantly to the cytosol, while its human ortholog (hUBE3a-Iso3) was enriched in the nucleus. The primate specific UBE3A isoform (hUBE3A-Iso2, <1% of total UBE3A) was found to be strictly cytosolic (1).

The importance of nuclear UBE3A for normal brain development is emphasized by the finding that selective abrogation of the nuclear, but not the cytosolic UBE3A isoform, causes behavioural phenotypes in mice (2). In addition, we found that a high percentage of AS-associated UBE3A missense variants, cause mistargeting of the short nuclear UBE3A isoform to the cytosol (3). These experiments highlight the importance of proper UBE3A localization and suggest that mistargeting of the nuclear isoform is sufficient to cause AS. Hence, we implemented the UBE3A localization assay as a first-tier screen to assess the pathogenicity of novel UBE3A missense variants (3).

Recently, three individuals diagnosed with AS have been described who carry the same genetic variant in *UBE3A*, which abrogates the start codon of hUBE3A-Iso1 (NM_130838.4:c.2T>C; (NP_570853.1):p.Met1Thr) and concomitantly results in a missense variant in both hUBE3A-Iso2 (NM_000462.5:c.71T>C; (NP_000453.2):p.(Met24Thr)) and hUBE3A-Iso3 (NM_130839.5:c.62T>C; NP_570854.1):p.(Met21Thr)). Despite complete loss of the most abundant nuclear hUBE3A-Iso1, these patients show higher developmental skills than typical AS patients (4), indicating that the remaining isoforms can compensate to some extent for the loss of nuclear hUBE3A-iso 1. However, the effect of this missense mutation on the localization of the two other isoforms is unknown. Given the importance of nuclear UBE3A for normal brain development and the fact that hUBE3A-Iso3 is the second most abundant isoform in neurons we assessed the effect of the p.Met21Thr variant on the localization of hUBE3A-Iso3.

Surprisingly, recombinant overexpression of hUBE3A-Iso3^p.Met21Thr^ in mouse primary hippocampal AS neurons and U2-OS cells revealed similar UBE3A labelling intensities in the nucleus and cytosol, suggesting hUBE3A-Iso3 localization is sensitive c (**Figure 1C**). Notably, in the previous study (1) a hUBE3A-Iso3^p.Met21Ala^ construct was used instead of hUBE3A-Iso3^WT^, in order to prevent translation from the second methionine (p.Met21), which could result in expression of hUBE3A-Iso1 protein, thereby hampering localization studies (**Figure 1B**).

**Figure 1.**
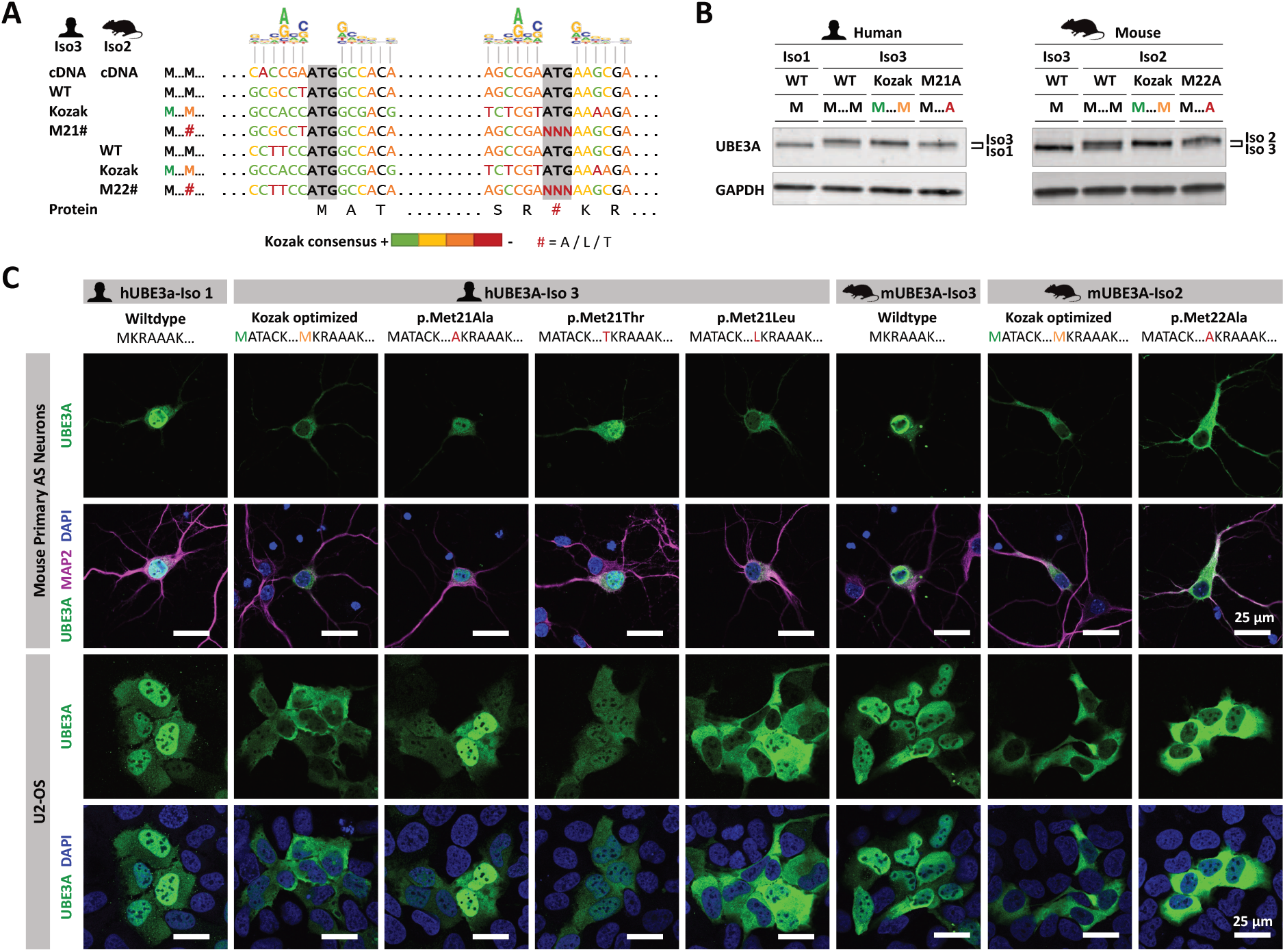
Localization of hUBE3a-Iso3 is sensitive to amino acid substitutions at the p.Met21 position. (**A**) Kozak sequences of the primary start codon and second methionine of hUBE3A-Iso3 and mUBE3A-Iso2 in the cDNA, WT vector, Kozak optimized vector, and vectors with p.Met21 or p.Met22 variants. Adherence to the Kozak consensus sequence is indicated with a colour scale. (**B**) Western blot analysis of UBE3A expression in U2OS cells recombinantly expressing hUBE3A-Iso1, hUBE3A-Iso3^WT^, hUBE3A-Iso3^KOZAK^, hUBE3A-Iso3^p.Met21Ala^ or their mouse orthologs mUBE3a-Iso3, mUBE3a-Iso2^WT^, mUBE3A-Iso2^KOZAK^, mUBE3A-Iso2^p.Met22Ala^. GAPDH was used as a loading control. (**C**) Immunofluorescence imaging of recombinantly expressed hUBE3a and mUBE3a isoforms and variants (untagged) in hippocampal mouse neurons (E16.5 Ube3a^m-/p+^ (AS) embryo’s, transfected at DIV7, fixed at DIV10, top) and U2-OS cells (fixed 48 hours post-transfection, bottom). UBE3a localization was visualized by anti-UBE3a staining (Alexa488, green), anti-MAP2 (Alexa647, magenta) staining was included as a neuronal marker and DAPI as nuclear counterstain. Scale bars: 25 μm. **Alt Text Figure 1A** Sequence alignments of the Kozak region surrounding the first and second methionine in human UBE3A Isoform 3 and mouse UBE3A Isoform 2. **Alt Text Figure 1B** A Western blot shows overexpression of UBE3A results in two bands, indicating translation from both methionines, while for Kozak optimized constructs the longer isoform is predominantly expressed. **Alt Text Figure 1C** Immunofluorescence microscopy images show predominant nuclear localization for recombinantly expressed hUBE3A-Iso1, hUBE3A-Iso3^p.Met21Ala^ and mUBE3A-Iso3, predominant cytosolic localization for hUBE3A-Iso3^p.Met21Ala,^ hUBE3A-Iso3^p.Met21Leu,^, mUBE3A-Iso2^KOZAK^, mUBE3A-Iso2^p.Met22Ala^ and equal distribution between nucleus and cytosol for hUBE3A-Iso3^p.Met21Thr^.

Realizing that hUBE3A-Iso3 is sensitive to amino acid changes at the p.Met21 position, we revisited the previous study and introduced silent mutations in the hUBE3A-Iso3^WT^ construct to optimize the Kozak sequences of the primary start codon (p.Met1), and more importantly, disfavour translation from the second methionine (p.Met21) (hUBE3A-Iso3^KOZAK^) (**Figure 1A**). Overexpression of hUBE3A-Iso3^KOZAK^ resulted in almost exclusive expression of the long isoform in U2-OS cells as assessed by Western blot (**Figure 1B**). Surprisingly, immunofluorescence staining revealed distinct cytosolic localization of hUBE3A-Iso3^KOZAK^ in U2-OS cells as well as in mouse primary hippocampal AS neurons, while recombinant expression of hUBE3A-Iso3^p.Met21Ala^ showed clear nuclear enrichment in both cell types (**Figure 1C**) as described previously (1). Thus, the previous conclusion that hUBE3A-Iso3 localizes to the nucleus, in contrast to its cytosolic mouse ortholog, is incorrect and resulted from the modification of the p.Met21 to which hUBE3A-Iso3 localization is highly sensitive.

This observation prompted us to also revisit the localization of the mouse long isoform (mUBE3A-Iso2), as the previous study also made use of a mUBE3A-Iso2^p.Met22Ala^ construct in which the second methionine (p.Met22) was substituted for an alanine. We used the same approach to generate a mUBE3A-Iso2^KOZAK^ construct for recombinant expression (**Figure 1A**), resulting in almost exclusive expression of the long isoform in U2-OS cells (**Figure 1B**). Upon overexpression of the mUBE3A-Iso2^KOZAK^ cytosolic localization of UBE3A was observed in U2-OS cells as well as in mouse primary hippocampal AS neurons (**Figure 1C**). While localization of hUBE3A-Iso3 is sensitive to changes on the p.Met21 position, mUBE3A-Iso2 is not for the equivalent p.Met22 position, and mUBE3a-Iso2 localization is predominantly cytosolic as previously reported (1,2).

Notably, to prevent translation from the second methionine in hUBE3A-Iso3 Chamberlain and co-workers introduced a leucine at position 21 (p.Met21Leu), an amino acid substitution that was predicted to have a neutral impact (5). Indeed, like hUBE3A-Iso3^KOZAK^, hUBE3A-Iso3^p.Met21Leu^ showed predominant cytosolic localization (**Figure 1C**).

Taken together, these data show that, like its mouse orthologue, the long hUBE3A-Iso3 localizes predominantly to the cytosol. The nuclear-cytosolic distribution of UBE3A in human and mouse is thus maintained by the orthologous UBE3A isoforms, in contrast to our previous findings. In addition, we show that the AS-associated p.Met21Thr variant causes a (partial) shift in the localization of hUBE3A-Iso3 to the nucleus. Consequently, individuals carrying the p.Met21Thr variant have both nuclear as well as cytosolic UBE3A, albeit at much lower levels. Hence, the milder phenotype of these individuals may be more reflective of lower UBE3A protein levels, rather than changes in UBE3A localization.

## Materials and Methods

### Cloning

Constructs for recombinant expression of hUBE3A^WT^ and mUBE3A^WT^ isoforms expressed under a CAG promotor were previously generated (1,2) with *Asc*I and *Not*I restriction sites flanking the *UBE3A* gene. *UBE3A* variants were generated through site directed mutagenesis, resulting in the N-terminal sequences as shown in **Table 1**. Sequences of all constructs were verified by Sanger sequencing (Macrogen) and plasmids were purified prior to use (Midi plasmid kit, QIAGEN).

**Table 1.**
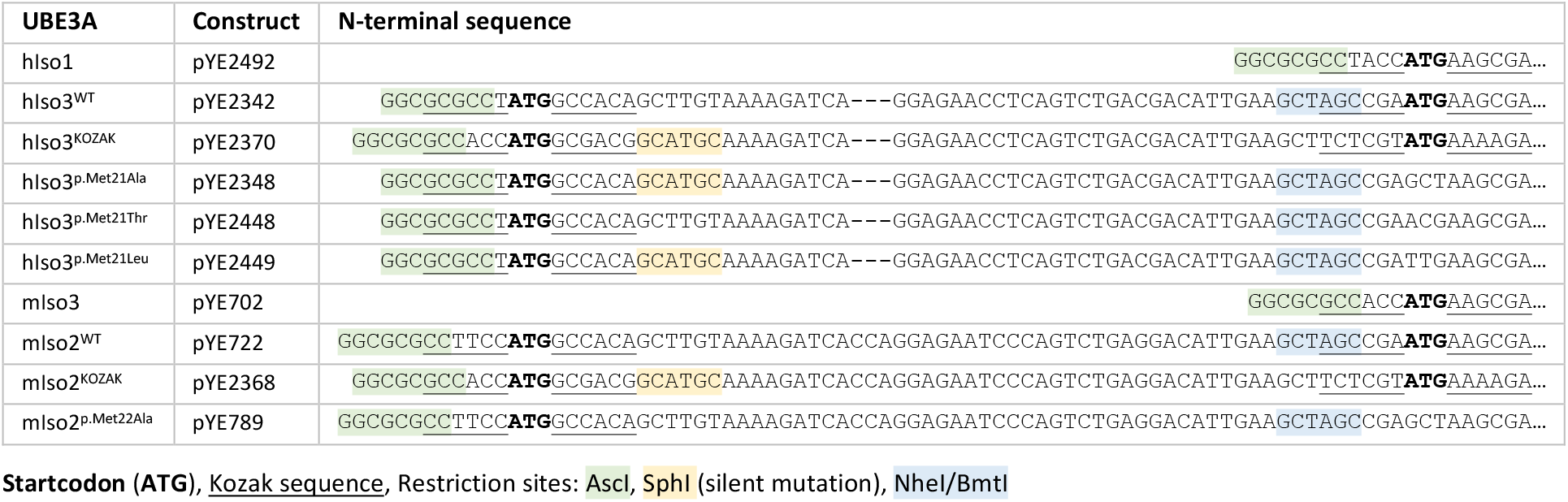
N-terminal sequences of UBE3A constructs.

### Cell Culture & Transfection

Mouse hippocampal primary AS neurons and U2-OS cells were cultured as previously described (1), with the following modifications. U2-OS cells were seeded in a 6--well plate containing a non-coated glass coverslip (300.000 cells/well) and transfected using 3 µg plasmid DNA and 9 µg polyethyleneimine per well. Coverslips were fixed 48 hours post -transfection for immunofluorescence, while remaining cells were harvested in 500 µL lysis buffer (50 mM Tris pH6.8, 2% SDS, 1x protease inhibitor (Roche Complete EDTA-free)) snap-frozen and stored at -70 °C until further processing.

### Immunofluorescence

Immunofluorescence was performed as previously described (1), with the following modifications. After staining with primary antibody (1:750 mouse-anti-UBE3a (SAB1404508, Sigma Aldrich) for U2-OS and neurons, 1:500 guinea pig-anti-MAP2 (188004, Synaptic Systems) for neurons) and secondary antibody (1:200 donkey-anti-mouse-Alexa Fluor™ 488 (715-545-150, Jackson Immunoresearch) for U2-OS and neurons, 1:200 donkey-anti-guinea pig-Alexa Fluor™ 647 (706-605-148, Jackson Immunoresearch) for neurons), coverslips were mounted on a drop of ProLong™ Gold antifade reagent with DAPI (Invitrogen). Confocal images were taken on a LSM700 Zeiss Confocal Laser Scanning Microscope using a 63x Oil objective.

### Western blot

Samples were thawed on ice and lysed by probe sonication (3 × 3 sec). Protein concentrations were determined by BCA assay (Thermo Scientific Scientific) and samples were diluted in lysis buffer (50 mM Tris pH 6.8, 2% SDS) and supplemented with 50 mM DTT and 1x XT Sample buffer (Bio-Rad) to a final protein concentration of 0.25 µg/µL. Samples were denatured (5 min, 95 °C) and 1.25 µg sample was resolved on a 4-12% acrylamide Criterion XT Bis-Tris gel (Bio-Rad) in MOPS buffer at 100 V for 20 min and subsequently at 150V for 190 min on ice along with PageRuler™ Plus Prestained Protein Ladder (Thermo Scientific. Proteins were transferred to a 0.2 µm nitrocellulose membrane by Trans-Blot Turbo™ Transfer system (Bio-Rad). Membranes were washed with TBS (50mM Tris pH 7.6, 150 mM NaCl) and blocked with 5% skim milk powder (Sigma Aldrich) in TBST (TBS with 0.05% Tween20) (1h, rt). Membranes were subsequently incubated with primary antibody mouse-anti-UBE3a (E8655, Sigma) (1:1000 in TBS-T, O/N, 4°C) and rabbit-anti-GAPDH (2118S, Cell Signaling Technologies) (1:2000 in TBS-T). After incubation blots were washed with TBS-T (3 × 5 min) and incubated with goat-anti-mouse (LI-COR Biosciences, IRDye 800CW, 926-32210) and goat-anti-rabbit (LI-COR Biosciences, IRDye 800CW,926-32211) (both 1:15.000 in TBS-T, 1 h, rt). After rinsing the membrane with TBS-T (3 × 5 min) and TBS (3 × 5 min) fluorescence was detected by scanning on the Odyssey CLx (LI-COR Biosciences).

## Acknowledgements

Ilse Wallaard for setting up the mouse primary neuron cultures.

## Author Contributions

MB and MF initiated the study. AE and RV performed the reported experiments. AE, YE, and BD wrote the manuscript. All authors reviewed and approved of the submitted manuscript.

